# Structural redundancy in supracellular actomyosin networks enables robust tissue folding

**DOI:** 10.1101/530816

**Authors:** Hannah G. Yevick, Pearson W. Miller, Jörn Dunkel, Adam C. Martin

## Abstract

Tissue morphogenesis is strikingly reproducible. Yet, how tissues are robustly sculpted, even under challenging conditions, is unknown. Here, we combined network analysis, experimental perturbations, and computational modeling to determine how network connectivity between hundreds of contractile cells on the ventral side of the *Drosophila* embryo ensures robust tissue folding. We identified two network properties that mechanically promote robustness. First, redundant supracellular cytoskeletal network paths ensure global connectivity, even with network degradation. By forming many more connections than are required, morphogenesis is not disrupted by local network damage, analogous to the way redundancy guarantees the large-scale function of vasculature and transportation networks. Second, directional stiffening of edges oriented orthogonal to the folding axis promotes furrow formation at lower contractility levels. Structural redundancy and directional network stiffening ensure robust tissue folding with proper orientation.

## Introduction

A hallmark of embryonic development is that it is reproducible. To achieve a stereotypic outcome, morphogenesis must be robust to genetic and environmental variations. Progress has been made in understanding how the embryo can use genetic or molecular strategies to overcome perturbations (Félix and Barkoulas, 2015; Whitacre, 2012). For example, redundancy in gene regulatory elements, such as shadow enhancers, promotes more robust gene regulation during development (Frankel et al., 2010; Hong et al., 2008; Perry et al., 2010). Redundancy in protein function or chaperone activity ensures reproducible physiological outcomes (Burga et al., 2011; Siegal and Rushlow, 2012; Zheng et al., 2013). Although these studies provided important insights into how reproducibility is regulated at the cell level, how reproducibility is achieved for populations of physically interacting cells is unknown.

Tissue contraction is a key mode of morphogenesis where force is propagated across distances that are far longer than the cell length-scale (Fernandez-Gonzalez et al., 2009; Galea et al., 2017; Hutson et al., 2003a; Martin et al., 2010; Varner and Taber, 2012). Tissue contraction can bring opposing cell sheets together, a mechanism used in both development and wound healing (Davidson et al., 2002; Kiehart et al., 2000). In addition, the selective contraction or expansion of one surface on an epithelial sheet (i.e., apical or basal) can result in tissue folding and cell invagination (Gutzman et al., 2008; Heer et al., 2017; Krueger et al., 2018; Polyakov et al., 2014; Sui et al., 2018). In several cases where there is tissue contraction, a supracellular actomyosin network is present across the field of cells (Hannezo et al., 2015; Martin et al., 2010; Nishimura et al., 2012; Roper, 2013; Skoglund et al., 2008). The function of supracellular actomyosin networks in morphogenesis is poorly understood. For example, do cells connect equally with all neighbors or can some cells take on more important roles in connecting parts of the tissue? How does the pattern of mechanical interactions in a tissue enable robust morphogenesis?

Ascertaining the network of cell interactions that enables a tissue to change shape is critical to understand morphogenesis. A limitation of past studies was the lack of methods to analyze both the cellular and supracellular structure of actomyosin networks. Here, we adapted an algorithm, originally developed to identify filamentous structures in the universe (Sousbie et al., 2011), to trace the filamentous structure of the supracellular actomyosin network during the folding of the *Drosophila* ventral furrow. *Drosophila* ventral furrow formation is an essential developmental step, which occurs when the embryo consists of a single layer of cells surrounding a central yolk. At this stage of development, ventral cells undergo apical constriction, which causes monolayer folding and internalizes the ventral cells (Leptin and Grunewald, 1990; Sweeton et al., 1991). The furrow or fold is reproducibly oriented along the anterior-posterior (a-p) axis, which is the direction of highest tension (Chanet et al., 2017; Martin et al., 2010). Analysis of wild-type and experimentally degraded networks coupled with computational modeling identified two mechanisms that make tissue folding robust.

First, redundancy in cytoskeletal connections creates multiple mechanical paths spanning the ventral tissue, which ensures at least one path across the tissue when there is local damage. Second, stiffening of network connections along the a-p axis enables furrow formation, even at lowered contractility levels.

## Results

### Ventral furrow formation is robust to tissue damage in the absence of wound healing

To investigate whether tissue folding is robust to cell/tissue damage, we ablated specific numbers of cells and determined whether the tissue could fold. We systematically injured embryos (4 – 32 cells) using high-intensity, two-photon laser excitation in cells prior to myosin II (myosin) accumulation and apical constriction. These targeted ablations created patches of damaged cells that did not apically constrict (Fig. 1A, C). In contrast to other stages in *Drosophila* development (Fernandez-Gonzalez and Zallen, 2013; Rodriguez-Diaz et al., 2008), wounding prior to apical constriction (i.e. late cellularization stage) did not result in an actomyosin purse-string or rapid wound closure (Fig. 1C, E, Supplemental Movie 1). Instead, the wound increased in area as surrounding cells began to recruit myosin and constrict. In addition, during the folding process, the wound anisotropy grew, suggesting that surrounding contractile cells stretched the wound (Fig. 1F). Tracking tissue flow towards the midline with PIV analysis demonstrated that damage resulted in slower movement, unlike cases where there is wound healing (Hutson et al., 2003b) (Fig. 1G). Despite wounding ventral cells in this manner, the tissue folded in all cases and successfully reached the larval stage (Fig. 1 B, D, n=5/5 embryos). Thus, *Drosophila* ventral furrow formation is robust to physical damage, even in the absence of a wound healing mechanism.

**Figure 1.**
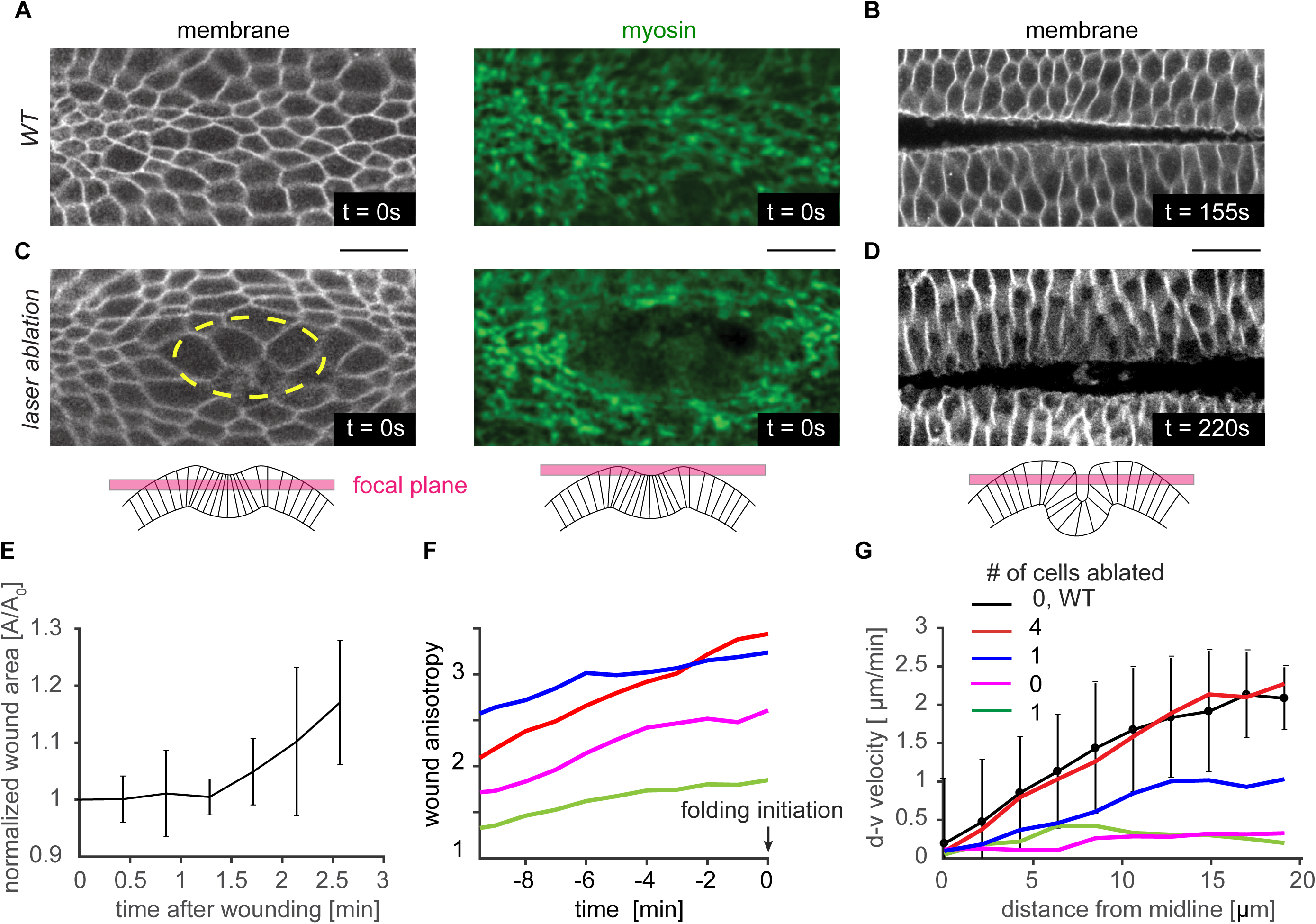
Ventral furrow folding is robust to tissue damage. (A) Cell outlines (plasma membrane marker) and the myosin supracellular network at *t=0*, the time of folding initiation, and (B) during tissue folding. (C) *t=0* for a tissue which was damaged via laser ablation. Cells were ablated prior to myosin accumulation in the region outlined in yellow. (D) The ventral tissue folds in the laser ablation condition as well. For (A-D) membrane is imaged subapically (using Gap43::mCherry), and apical myosin is visualized in a Z-projection (using sqh::GFP). (E) Ablation after wounding does not undergo rapid wound healing resulting in closure. Rather the hole opens larger as surrounding cells begin to recruit myosin. For this plot only *t=0* is time of ablation. N=3 embryos. (F) The wound stretches along the anterior-posterior axis (anisotropy increases) prior to folding initiation. Embryos are aligned to *t=0*. (G) The velocity of the cells towards the midline at the time of folding (*t=0*) slows with increased tissue damage, but all embryos still fold. (F,G). N=1 embryo for each of the 4 wound sizes. Scale bars: 20 µm.

### Supracellular ‘mechanical paths’ connect hundreds of cells in the ventral furrow

The striking robustness of the folding process led us to investigate mechanisms that could produce this mechanical resilience. We hypothesized that patterns of linkages between individual force-generating cells establishes robust folding. In the *Drosophila* ventral furrow, mechanical linkages are associated with supracellular, apical actomyosin fibers that connect between cells through spot adherens junctions, in an end-on manner (Martin et al., 2010) (Fig. 2A, Supplemental Movie 2). Apical myosin fibers colocalize with apical actin filaments (F-actin) and these fibers recoil when they are cut, suggesting they support tension across a cell’s apical surface (Martin et al., 2010). Thus, supracellular myosin structures represent ‘mechanical paths’ that form a network connecting across cells in the tissue.

**Figure 2.**
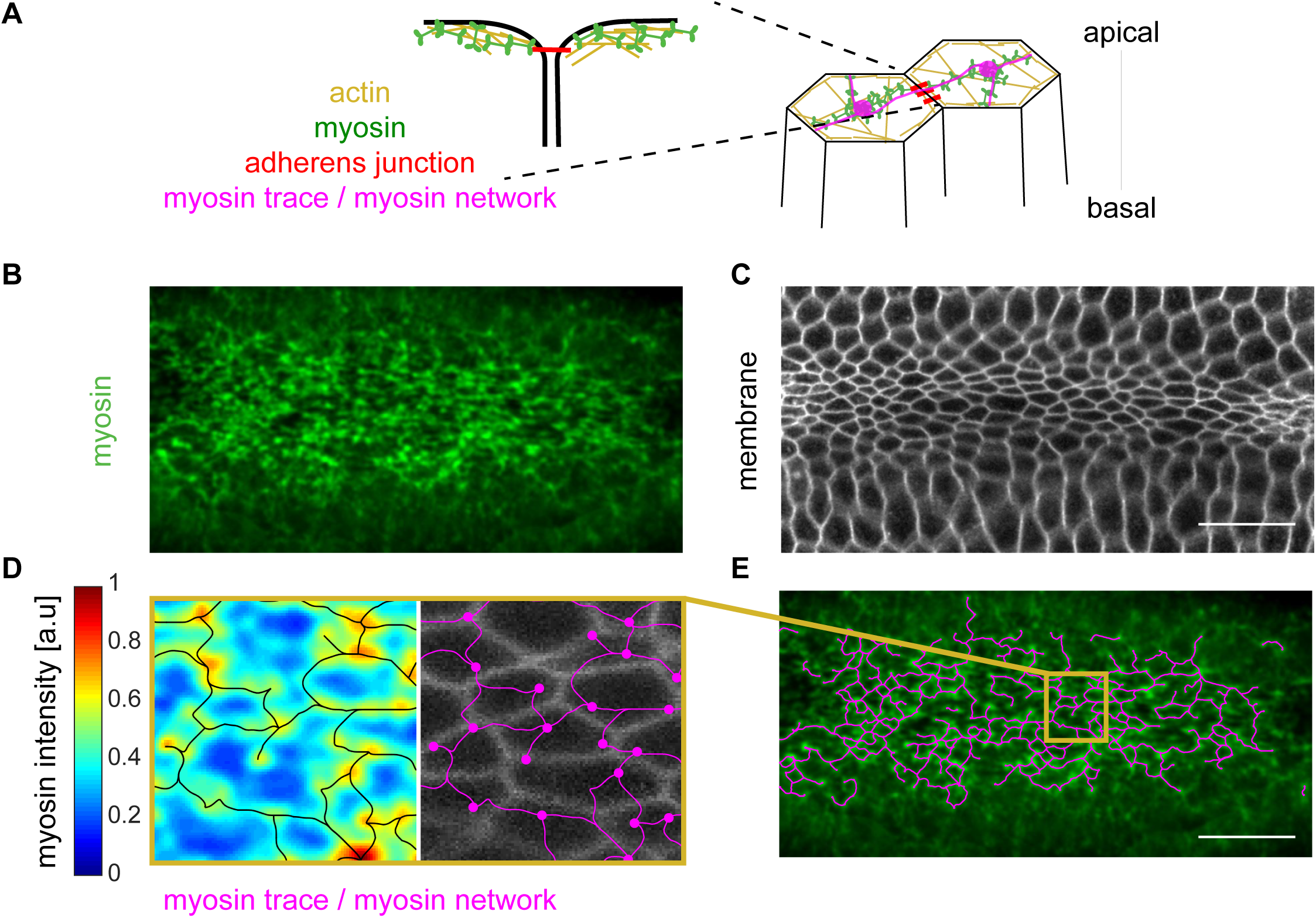
Developing a method to trace the supracellular actomyosin network. (A) Myosin is predominantly activated apically. It interacts with actin to form actomyosin structures across the apical surface of cells. These fibers connect between cells in an end-on manner at spot adherens junctions. (B) Z-projection of myosin (green, sqh::GFP), and (C) subapical slice of membrane (Gap43::mCherry) channel at the time of folding initiation. Myosin forms a network across the entire apical surface of the ventral tissue just prior to folding. (D) Apical myosin intensity (blue-red heat map) was traced using the *DisPerSE* algorithm (Sousbie, 2011). The trace follows the topology of the intensity signal and forms a network (magenta) where nodes correspond to local maxima and edges correspond to the topological ridges between these peaks. The myosin network (magenta) does not correspond to the cell membranes (grayscale). (E) The trace of the supracellular myosin network (magenta) across hundreds of cells in the ventral tissue. Scale bars: 20 µm.

Evidence exists that some minimal connectivity of this network is needed for folding. For example, disrupting the intercellular junctions disrupts folding (Dawes-Hoang et al., 2005; Martin et al., 2010; Sawyer et al., 2009). In addition, laser ablations that sever connectivity by isolating cells along the anterior-posterior axis disrupts folding (Chanet et al., 2017). However, whereas individual supracellular myosin connections have been described at the cell level (Martin et al., 2010), the global network and the importance of its structure have not been determined.

Supracellular actomyosin networks have not been rigorously analyzed due to the difficulty of defining subcellular structure in noisy tissue-scale images (Fig. 2B, C). Therefore, to determine this global network structure, we adapted a topological algorithm to trace the filamentous supracellular myosin network in noisy confocal images (Fig. 2D, E). This framework, called *Discrete Persistent Structures Extractor* (*DisPerSE*), was originally developed to trace the filamentous structures in noisy Astrophysics data (Sousbie et al., 2011). *DisPerSE* employs Morse-Smale theory to partition scalar fields based on the gradient of the input image and a persistence extractor to smooth the signal (Supplementary Fig. 1, Methods). We extracted the peaks of myosin intensity (which became the network “nodes”) and ridge-lines connecting the nodes (which became the network “edges”). Visually, the trace of the myosin network in wild-type embryos corresponded closely with myosin structure and clearly spanned the entire ventral furrow (Fig. 2D, E).

To confirm that the traced myosin structure captures intercellular connectivity, we compared the wild-type myosin network to the myosin network in embryos depleted of an essential adherens junction component, α-catenin. As expected, α-catenin depletion disrupted supracellular myosin network formation; instead, myosin coalesced into isolated nodes in the middle of each cell apex (Martin et al., 2010) (Supplementary Fig. 2A). The tracing algorithm correctly showed that supracellular myosin fibers were no longer present in α-catenin-depleted embryos and that the network length was significantly reduced (Supplementary Fig. 2B). Therefore, our decomposition of the apical myosin network into nodes and edges, captures the intercellular connectivity of cytoskeletal structures across the tissue.

### The supracellular actomyosin network grows dynamically during folding

Using this topological structure identification algorithm to map the entire supracellular myosin network, we first determined how the supracellular myosin network is assembled during folding. For this study, we aligned embryos based on the time of tissue folding. We define this time, *t=0*, as the time when out-of-plane deformation is first perceived.

For all embryos analyzed, the number of nodes in the myosin network increased linearly with time (Fig. 3A). However, nodes were not added everywhere in the ventral region with equal probability. Nodes accumulated fastest along the ventral midline, with the gradient in node density along the dorsal-ventral axis growing over time (Fig. 3B). A dorsal-ventral gradient in the probability of having a myosin node is consistent with there being a multicellular gradient in average apical myosin activity (Heer et al., 2017). While the network increased in size, individual myosin nodes existed transiently, exhibiting a mean lifetime of 100-150 seconds (Fig. 3C and Supplementary Fig. 2C).

**Figure 3.**
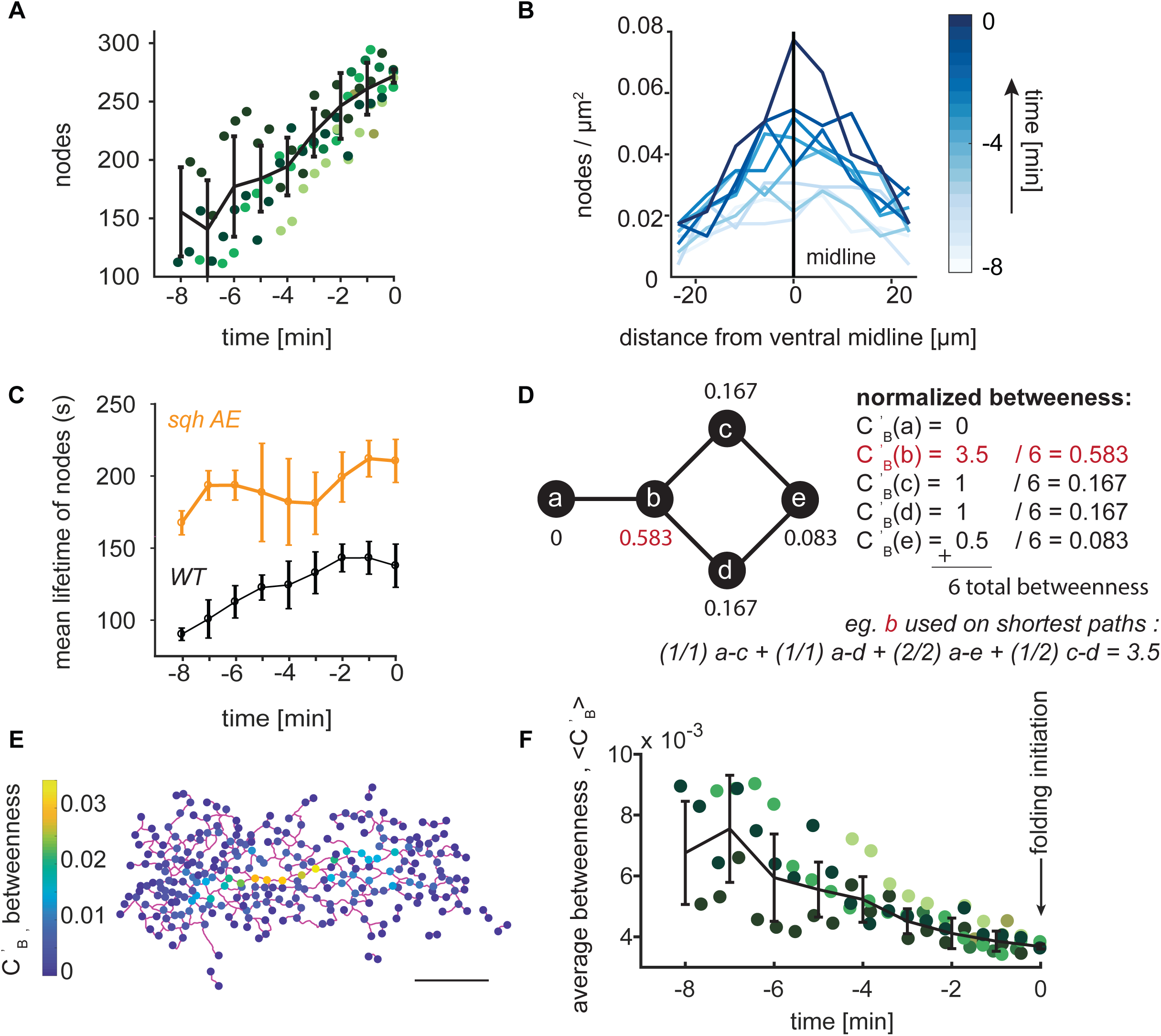
Nodes in the supracellular myosin network become increasingly redundant during ventral furrow formation. (A) The number of network nodes grows leading up to folding (*t=0*). Plotted is the number of nodes detected in an embryo as a function of developmental time; the different colors represent different embryos. Trend line (black) represents mean ± s.d. (B) Node density is present in a ventral-dorsal gradient. The highest node density is at the ventral midline (0 on x-axis) with node density decreasing away from the ventral midline. The gradient increases in magnitude leading up to folding. (C) Node lifetime depends on myosin regulation. Plot shows the mean ± s.d. of node lifetimes for wild-type or *sqh*-AE mutant tracked nodes. Pulsed cycles myosin accumulation and disassembly normally observed are abolished in the *sqh*-AE mutant. (D) An example network and the measure of normalized betweeness centrality, C_b_^’^, which is calculated for each node. Node *b* has the highest normalized *betweeness* as it the most important node in connecting other sets of nodes. (E) Representative wild-type supracellular network where *betweenness* was calculated for each node (color bar). Network edges connecting the nodes are in magenta. (F) The average *betweenness* for each node decreases over time leading up to the time of folding initiation (*t=0*). Trend line (black) represents mean ± s.d. (A,F) Each color represents a different embryo. N=6 embryos. (B,E) data is from a single representative embryo. (C) N=3 embryos for each condition. Scale bar: 20 µm.

To determine if these node dynamics reflected myosin turnover, we examined node lifetime in a myosin light chain mutant, the *sqh-AE* mutant (Vasquez et al., 2014). The *sqh-AE* mutant disrupts myosin turnover, likely because it uncouples myosin regulation from dynamic upstream signals and because it compromises motor activity (Vasquez et al., 2016, Mason et al., 2016). Myosin node lifetime was significantly increased in *sqh-AE* mutant embryos, suggesting that the myosin node dynamics we observed in the network are due to cytoskeletal and signaling dynamics (Fig 3C and Supplementary Figure 2C,D). In summary, the supracellular myosin network maintains global connectivity while individual network nodes and, therefore, their corresponding connections are dynamic.

### Supracellular ‘mechanical paths’ are redundant in the wild-type actomyosin network

To determine how the global network connectivity couples distinct regions of the tissue, we sought to measure the relative importance of nodes in connecting the network. To identify nodes that heavily support the overall network connectivity, studies of network resilience in subway systems (Derrible, 2012), airports (Guimerà et al., 2005), and roads (Kermanshah and Derrible, 2017; Lämmer et al., 2006) have used the *betweenness centrality, C*_*B*_. *Betweenness centrality, C*_*B*_, measures the fraction of times a node is on the shortest path between any other two nodes (Freeman, 1978). To ensure our calculated *betweenness* did not simply reflect network size, we normalized the measurement to the total number of paths, 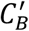 (Fig. 3D) (Methods). Therefore, nodes with higher *betweenness* are more critical in connecting regions of the tissue.

We measured the normalized *betweenness* of all nodes in the network of 5 wild-type embryos (Fig. 3E, Supplemental Fig. 3) at each developmental time. For all embryos, the mean node *betweenness* decreased to a consistently low value at which point the tissue folded (Fig. 3F). Therefore, individual nodes in the wild-type network become increasingly redundant (i.e., less important in connecting parts of the tissue) leading up to folding.

### Network degradation decreases redundancy, but does not prevent folding

To test the function of redundancy in folding, we degraded the network in two different ways and assessed network structure. First, we examined the network structure of laser-ablated embryos (Fig. 1C). Second, we overexpressed a constitutively active form of the myosin binding subunit (MBS^N300^) of myosin phosphatase to downregulate myosin activity (Fig. 4A). In contrast to laser ablation, MBS^N300^ overexpression resulted in a more uniform degradation of the supracellular myosin network; clusters of cells failed to accumulate apical myosin and constrict, similar to mutants that disrupt signaling upstream of RhoA (Fig. 4B) (Costa et al., 1994; Parks and Wieschaus, 1991; Xie et al., 2016). Myosin degradation reduced, but did not abolish, the gradient in node density along the ventral-dorsal axis (Supplementary Fig. 4A). Similar to our results with laser ablation (Fig. 1 C, D), folding was observed even when MBS^CA^ expression caused a 65% reduction in the number of nodes across the tissue (Supplemental Movie 3). Thus, our results with laser ablation and MBS^N300^ expression demonstrated that tissue folding is robust to network degradation.

**Figure 4.**
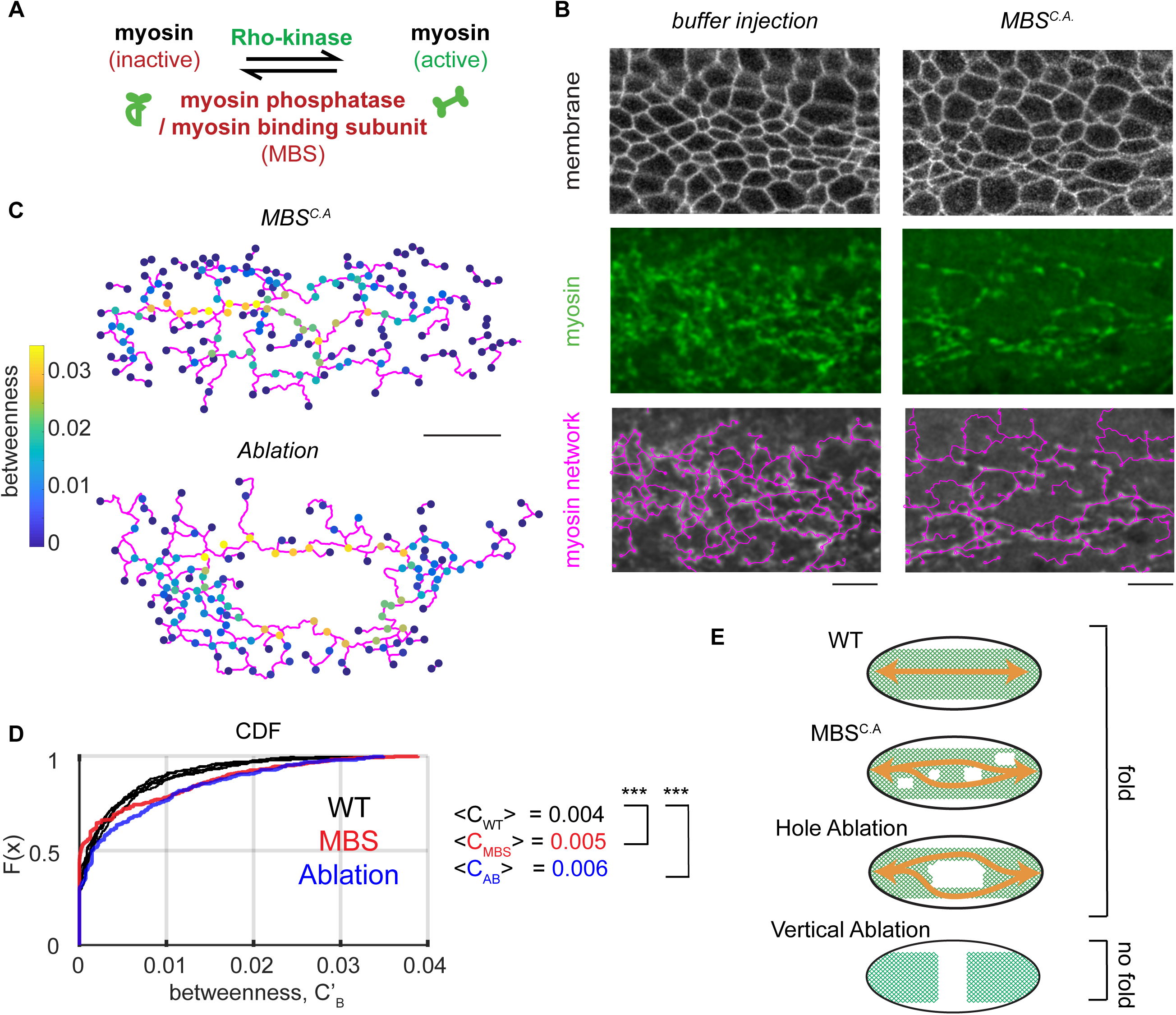
Decreasing network redundancy does not prevent folding. (A) Cartoon showing myosin regulation. Rho-kinase activates myosin by phosphorylating its regulatory light chain and also by inhibiting myosin phosphatase. Myosin phosphatase/MBS inactivates myosin. (B) Constitutively active MBS^N300^ (MBS^CA^) degrades the supracellular myosin network. Images show subapical membrane (Gap43::mCherry, grayscale), Z-projected myosin (Sqh::GFP, green), and the network trace (magenta). (C) The node *betweenness* for representative embryos at the time of folding where 10 cells were ablated or the embryo was injected with MBS^N300^ mRNA. (D) The cumulative distribution of function of node *betweenness* in wild-type embryos (black) or a representative MBS^CA^–expressing embryo (red) and an embryo where 10 cells were ablated (blue). Note that laser ablation and MBS^CA^ shift the CDF curve down, which indicates a greater proportion of nodes with high *betweenness*. N = 6 WT embryos, 1678 nodes, Ablation N=2, 343 nodes. p value comparing WT to AB = 8×10^−5^. MBS N=2 embryos 401 nodes. p value comparing WT to MBS 3×10^−8^ (two-sample KS test). (E) Cartoon showing how redundant connections can continue to link (orange arrows) regions of the tissue after either myosin degradation (MBS^CA^) or laser ablation. Vertical laser ablations, however, remove all paths across the embryo. Scale bars (B): 10 µm, (D) 20 µm.

Degradation of the supracellular actomyosin network increased the relative importance of individual network nodes in connecting the tissue, but did not completely disconnect components of the network (Fig. 4C). The cumulative distribution function of *betweenness* values in wild-type, laser ablation, and MBS^CA^ overexpression conditions at the time of folding demonstrated that nodes in the network have higher normalized *betweenness* values upon network degradation (Fig. 4C, D). These results suggested that many more paths are present in the wild-type supracellular myosin network than are needed to fold the tissue. Network redundancy in the supracellular actomyosin network ensures a minimal path across the tissue, even when there is damage (Fig. 4E). Thus, tissues folded when degraded by either laser ablation or MBS^CA^ overexpression, unless the entire width of the ventral furrow domain was severed (Fig. 4E, bottom) (Chanet et al., 2017).

### Wild-type supracellular networks exhibit connectivity below the isostatic limit

Because we showed that the ventral furrow folds even with decreased network connectivity, we investigated what properties of the remaining connections enabled reproducible folding. We hypothesized that rigidity in the network could promote collective deformation of the tissue. Maxwell first described the *isostatic point* (Maxwell, 1864), which is the minimal average node degree in a network needed to achieve mechanical rigidity (i.e., infinite stiffness) (Fig. 5A). Below the isostatic point, ‘floppy modes’ exist, which are collective degrees of freedom that are not subject to a restoring force (Fig. 5B). Adding sufficient connections between nodes, as is the case for a truss in a bridge, rigidifies the structure.

**Figure 5.**
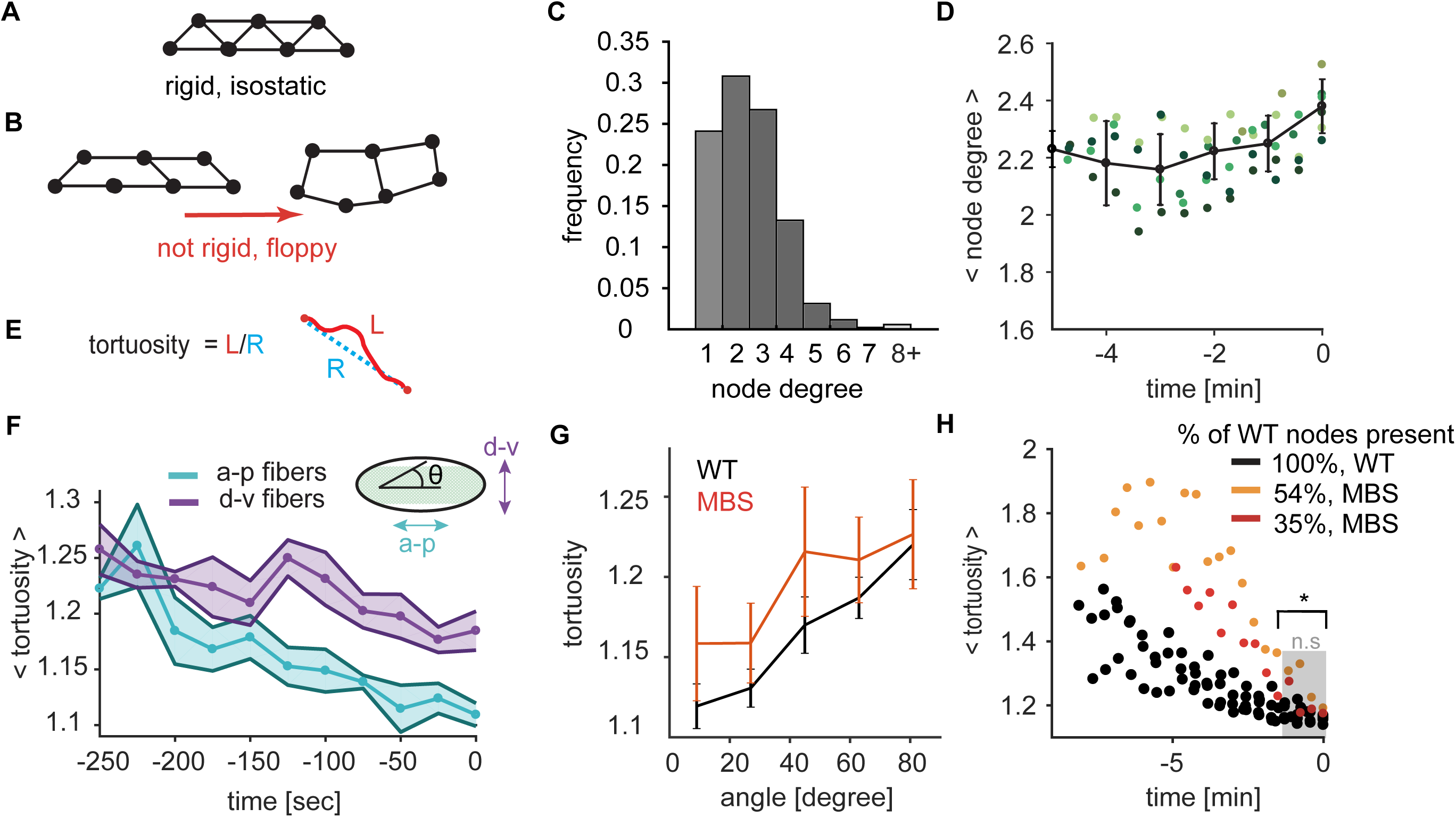
Directional network strain triggers folding. (A-B) Network rigidity is determined by the isostatic limit where the degrees of freedom are balanced by the constraints. The sample network of 7 connected nodes is rigid and cannot be deformed without straining the edges (A). Removing edges from the network in brings it below the isostatic limit and the network can be deformed with no restoring force (B). (C) The degree distribution of the nodes for wild-type networks at the time of folding. The degree of a node degree is defined as the number of edges which connect to it. N=6 embryos, 1361 nodes. (D) Average node degree is sub-isostatic and does not significantly increase during folding. Plotted is the mean ± s.d. node degree as a function of developmental time up to folding time. (E) Tortuosity is defined as the ratio of the arc length of an edge (L, red) to the distance between the two end points (R, blue). (F) Edges straighten preferentially along the a-p embryonic axis. Plotted bold line and data points show mean tortuosity as a function of time. Shaded region represents ± s.d. N=6 embryos, between 818 (t = - 250) to 1282 (t = 0) edges. Anterior-posterior edges are oriented from 0 – 18 degrees with respect to the midline and dorsal-ventral edges are between 71 – 90 degrees. (G) The degraded myosin network in MBS^CA^ expressing embryos also exhibits preferential straightening along the a-p axis (angle = 0). N=6 WT embryos, 1972 degrees, N=4 MBS embryos, 777 degrees. (H) Folding initiates for both wild-type and MBS^CA^ embryos when the edge connections straighten (i.e. edge *tortuosity* ~1.2). N=6 embryos (WT), 4 embryos (MBS), 1984 edges (WT), 777 edges MBS (at t = 0). The distribution between WT and MBS is statistically significant up until - 100 seconds before folding (t = −100, p = 0.01).

To determine whether wild-type supracellular myosin networks achieve isostacity, we measured the *degree* of each node across the network, which represents the number of edges emanating from it. For passive networks formed from spring-like forces between nodes, a rigid structure is achieved in two dimensions in the large network limit when the average node degree is 〈*z*〉 ≥ 4 (Maxwell, 1864). The average node degree in the wild-type supracellular actomyosin networks was low at the time of folding (〈*z*〉 = 2.38 ± 0.2, Fig. 5C, D), and it did not increase during the folding process. Thus, the density of edges in the supracellular actomyosin network does not reach the isostatic point, suggesting that connectivity controlled rigidity does not explain how the tissue folds.

### Folding is associated with the stiffening of a-p oriented connections

Studies of filament networks have shown that subisostatic networks can stiffen under strain (Sharma et al., 2016). A signature of a network under strain is edge straightening, which we quantified using a measure called *tortuosity*. The *tortuosity* of an edge is the ratio of the arc length of the edge to the distance between interconnected nodes (Fig. 5E). A value of 1 corresponds to an edge that is perfectly straight. We found that folding was associated with a preferential straightening of anterior-posterior (a-p) oriented edges. While a-p and dorsal-ventral (d-v) edges began with the same tortuosity, over time edges oriented along the a-p axis (angle 0 – 30°) decreased their *tortuosity* more significantly than those oriented in the d-v direction (Fig. 5F). This bias occurred in both wild-type and MBS^CA^ embryos (Fig 5G, Supplementary Fig. 4B-E). The straightening of a-p edges is consistent with the higher level of tissue tension along this axis (Chanet et al., 2017). Our finding shows that this tissue-level property is triggered by connections between cells becoming preferentially taut along the a-p axis.

To determine the relationship between network strain and folding, we measured mean *tortuosity* as a function of time for all network edges. In the wild-type embryos, *tortuosity* decreased prior to folding initiation (Fig. 5H). Interestingly, despite lower myosin levels, folding for the MBS^CA^ overexpressing embryos occurred at the same mean *tortuosity* for the myosin network as the edges in wild-type networks (Fig. 5H, WT and MBS^CA^ compared by two-sample KS test, p = 0.2), suggesting that the increase in a-p stiffness triggers tissue folding. Therefore, folding is not associated with increasing network connectivity, but is instead closely associated with the onset of directional network stiffening.

### Stiff a-p edges promote robust folding at low myosin levels

The directional bias in taut fibers along the a-p direction does not result in preferential area constriction in the a-p direction. Instead constriction occurs predominantly in the d-v direction (Heer et al., 2017; Martin et al., 2010). Therefore, to test how directionally biased (a-p) stiffness in a supracellular actomyosin network promotes d-v bending of the embryonic tissue, we employed a 3D continuum elastic shell model of the *Drosophila* embryo. Folding of the model embryo was driven by a gradient of myosin contractility applied to the shell surface, which is present in the embryo (Fig. 6A) (Heer et al., 2017). Myosin contractility induces folding by changing the local preferred curvature of the shell (Heer et al., 2017) (Methods). In addition to the myosin gradient, we modeled the supracellular actomyosin network by imposing a network of stiff elements on the surface. Because the edges of the network represent tense fibers connecting discrete points, we represented the mechanical effect of an edge by imposing an additional cost for the tissue to bend along the edge’s length (Fig. 6A). To make a general theoretical prediction as to how the network influences tissue folding, we made the network 10x stiffer than the background tissue. The trends in our model were confirmed by also implementing it with stiffnesses that were 4x and 8x times the background.

**Figure 6.**
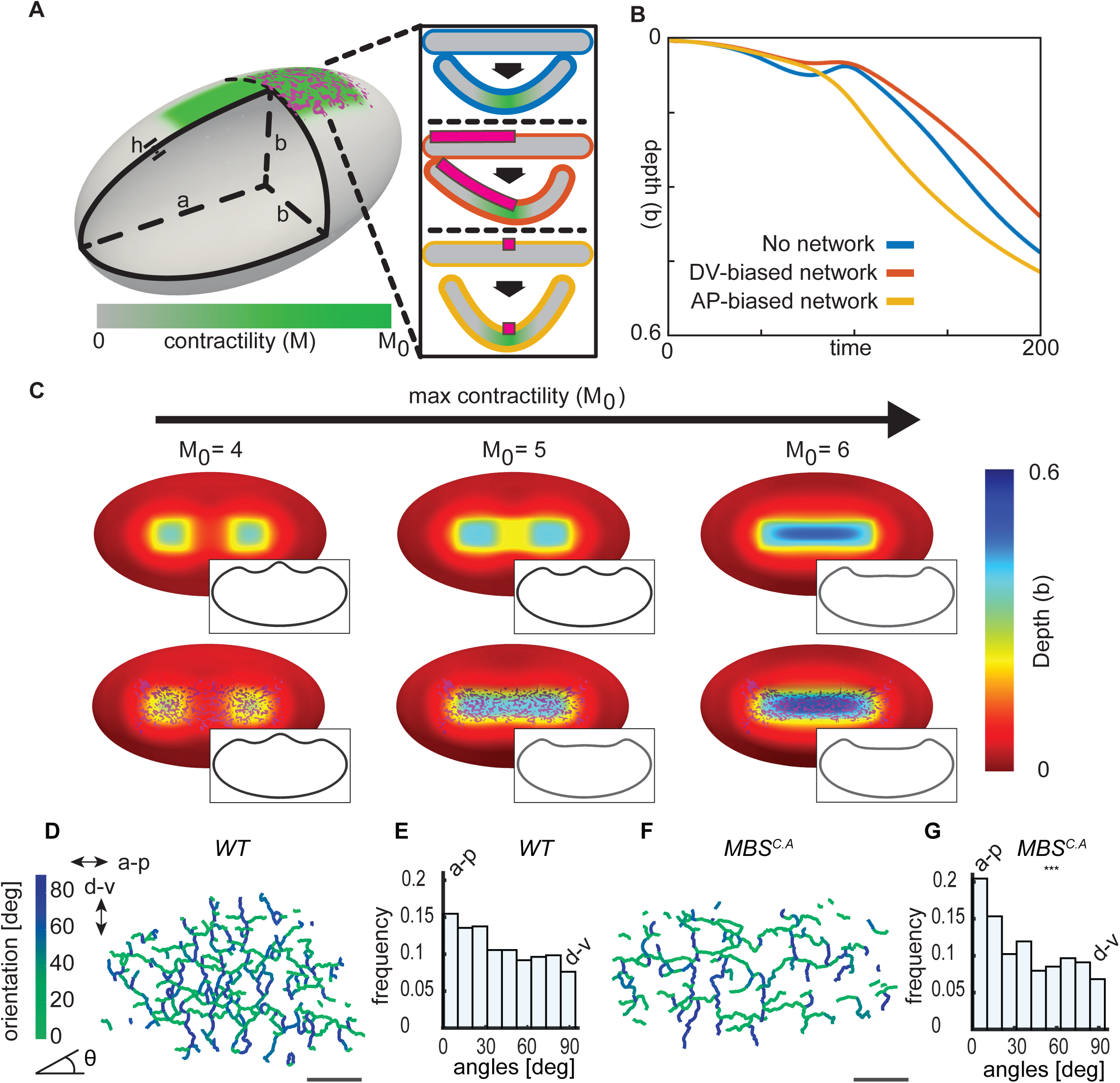
Directional network stiffness promotes robust furrow formation. (A) To model ventral furrow formation, a stiff myosin network (magenta) was superimposed on an elastic shell (grey), which contained a myosin contractility gradient (green) on one side. The insets show the effect of different fiber orientations (d-v, orange; a-p, yellow) on preferred curvature in the model. (B) A-p oriented stiff fibers accelerate folding *in silico*. Starting from an unfolded configuration, *M*_0_was slowly increased from 0 to 15, triggering folding. The depth of the furrow at the midpoint of the a-p axis was compared for three types of simulations: without a network (blue), with a network containing edges preferentially directed parallel to the a-p axis (yellow), and containing edges preferentially parallel to the d-v axis (orange). (C) A stiff a-p network promotes furrowing at lower contractility. Contractility was varied on elastic shells in the presence or absence of a stiff network, which resembled the experimentally measured network. Insets show mid-sagittal planes through the shells, indicating the lack of (red) or presence of (blue) a furrow. (D) Edges in a representative wild-type network at *t=0* are color-coded using their orientation with respect to the midline. (E) The wild-type supracellular myosin network has a slight bias in connections along the a-p axis. N = 6 embryos, 1935 edges. (F) Edges in a representative MBS^CA^ network at *t=0* are color-coded using their orientation with respect to the midline. (G) Myosin degradation results in a preferential loss of d-v connections, which increases the edge bias to the a-p direction. N = 4 embryos, 625 edges. p value 1.5×10^−12^ when compared to WT distribution (two-sample KS test). Scale bars: 20 µm.

First, we explored how the presence of a network influences folding. Starting with a level of myosin contractility that was just enough to generate a furrow, we simulated folding events with lower contractility levels and determined the effect of having a rigid network. For low myosin/contractility levels, rather than forming a furrow, depressions formed and the end of the contractile domain (Fig. 6C). However, imposing a rigid network resulted in furrow formation at contractility levels that normally would not result in a furrow. Therefore, the presence of a stiff network decreases the amount of contractility needed to form a furrow, making it more robust.

To explore how the directionality of network connections influences folding, we compared folding without a stiff network, to networks with stiff edges exhibiting preferred orientations along either the a-p or d-v axis. For a given network size, we found that folding speeds were the highest when network edges were preferentially oriented along the long axis of the shell when compared to the case where there was only contractility and no structural network (Fig. 6B, yellow vs. blue line). In contrast, networks where edges were preferentially oriented along the d-v direction folded slower than both the wild-type network and the a-p biased network (Fig. 6B, orange line). This result suggested that the structural properties of the supracellular actomyosin network promotes robust tissue folding and that the a-p connections in the network are most important for this effect.

Because the model predicted an important role of a-p oriented edges, we investigated whether there was a bias to network edges in embryos. In wild-type embryos, the myosin edges/connections were slightly biased toward alignment along the a-p axis (Fig. 6D-E). We then examined this bias in embryos overexpressing MBS^CA^. MBS^CA^ decreases myosin levels and the number of myosin nodes, but the remaining nodes were still distributed in a gradient from the ventral midline (Supplementary Fig. 4A). Interestingly, MBS^CA^ exhibited a greater bias in edge orientation (p value 1.5×10^−12^ two-sample KS test), preferentially retaining a-p edges (Fig. 6F-G). It is unclear why d-v edges are preferentially degraded upon MBS^CA^ expression, however, one possibility is that the greater tension/straightness of these edges makes them more stable (Fernandez-Gonzalez et al., 2009). Our model predicted that stiff edges oriented along the furrow axis increases folding efficiency and enables folding at lower motor activity. Our analysis of the network in MBS^CA^ overexpressing embryos highlights the importance of a-p network connections. MBS^CA^ embryos exhibited folding while predominantly having a-p connections, rather than d-v connections. Because contraction must happen in d-v for cells to obtain wedge shapes needed for the furrow shape, our results support a structural role for the supracellular actomyosin network, such as by providing directional tissue stiffness.

## Discussion

In this study, we developed a strategy to trace supracellular actomyosin networks involved in tissue morphogenesis. This strategy will be generally applicable to further investigations of both collective behavior in tissues and network structure in individual cells. Using this new approach, we discovered: 1) Many more supracellular connections are present in wild-type supracellular actomyosin networks than are minimally needed to fold the tissue. This network redundancy makes tissue folding robust. 2) The supracellular network plays a passive, structural role in folding with stiff a-p edges enhancing robustness. The network creates a ‘frame’ that determines final tissue shape; being taut/stiff along one axis and flexible along the orthogonal axis.

### Redundancy is a general strategy for network resilience

Our analysis of the supracellular network revealed a striking feature of the supracellular actomyosin network – individual nodes in the network are not important to connect different parts of the tissue. We also demonstrated this redundancy experimentally using both laser ablation and MBS^CA^ overexpression. In both cases, we could significantly degrade the network and tissue folding still occurred, arguing for redundancy in network connections. The presence of redundant paths across the tissue secured alternative spanning routes even upon the removal of some regions of the network. Therefore, the tissue can recover from damage and fold without wound healing.

Redundancy of tissue scale connections is a novel mechanism to mechanically ensure robustness during embryonic development. Embryos employ a similar strategy to that used in many other types of networks that exhibit large-scale function upon loss of connections due to attack or random failure. For example, seminal work in the field of network science on network resilience carried out by the US Airforce described the use of redundancy as a means of building communication systems that can withstand heavy nuclear attacks (Baran, 1964). Well-constructed transportation networks incorporate redundancy to optimize transport through the network under local fluctuations in transport capabilities (Corson, 2010). The cerebral cortex relies on a highly-interconnected network of communicating arterioles which can reroute flow to mitigate the effects of a single obstruction (Schaffer et al., 2006). Interestingly, while the myosin network in the ventral furrow is redundant, it is far from the optimal connectivity needed to maximize robustness under damage. Unlike the ventral furrow, scale-free networks, which have a power-law distribution of node degree (for example the internet), are more resistant to being broken into two disconnected parts upon node removal. We speculate that the relatively low average node degree of the supracellular myosin network has optimized the need for folding robustness relative to the energetic cost of forming a large number of connections per node.

One challenge in our study was to measure the exact network degradation threshold past which folding does not occur. Percolation theory predicts that removing a small number of nodes has only a limited impact on the network’s integrity with a sudden network failure at a critical threshold (Barabási, 2016). We hypothesize that the existence of such a critical threshold, made it difficult to experimentally identify the minimal connectivity needed for folding using MBS^N300^ overexpression. We either observed a network with reduced, yet less redundant a-p paths, or severe cellularization phenotypes and no network or folding. However, fully removing all a-p paths by severing the entire width of the constricting ventral domain via laser ablation did abolish folding (Chanet et al., 2017).

### The supracellular actomyosin network acts as a structural element that guides folding

This work defines a new role for supracellular actomyosin networks by demonstrating that they can play a structural role (like a frame) that promotes robust tissue morphogenesis. The presence of stiff network edges on the embryo surface, especially a-p oriented edges, mechanically promotes furrow formation by changing the local bending energy of the tissue. We showed that edges preferentially straighten when oriented along the direction of the future furrow and that there are more network connections oriented along this axis. We modeled these straight paths as stiff passive elements on an ellipsoid shell. Our physical model identified that these oriented connections promote furrowing along the correct axis at lower motor activity than is normally observed without the network. We also showed that a-p oriented edges are preferentially retained in folding tissues under myosin degradation. The directionality of these edges is orthogonal to the axis of preferential constriction, suggesting they do not simply mediate contraction. Therefore, tissue-scale actomyosin assemblies can have a structural function in sculpting tissues, in addition to actively generating the force needed to constrict cells. This ability of actomyosin to serve a structural role is reminiscent of supracellular actomyosin assembled at cell boundaries in some developing tissues (Monier et al., 2009; Umetsu et al., 2014). In these instances, tension directed along the boundary prevents the mixing of opposing groups of cells but does not promote cell or junction constriction. However, these tissue-scale actomyosin assemblies act solely in-plane and do not trigger the 3D shape change.

### Encoded robustness in the absence of wound healing

We report that robustness is encoded into morphogenetic movements even in the absence of a wound healing response, which is likely to be important for embryonic stages where such a response is not present. Wound healing is well-described mechanism for tissue resiliency (Redd et al., 2004). During dorsal closure, for example, a purse string at the margin of two sheets of epidermis pulls and seals together the tissue (Kiehart et al., 2017). When the actomyosin purse string is damaged, cells assemble a secondary cable at the injury site via wound healing (Rodriguez-Diaz et al., 2008). While wound healing can keep an embryo on the correct developmental program, mechanical robustness encoded into the tissue can allow a tissue to adjust immediately to damage. Further study is needed to understand how inherent robustness mechanisms interact with wound healing to promote reproducibility in embryonic development. Exploring inherent mechanisms of robustness in other tissues undergoing morphogenesis has the potential to elucidate new ways to control and reprogram tissues when both treating disease and when striving to engineer reproducible tissue shape *in vitro*.

## Acknowledgments

We would like to thank current and former members of the Martin lab for discussion and advice, most notably Claudia Vasquez and Mike Tworoger. We would also like to thank Norbert Stoop and Aden Forrow for helping to initiate the data analysis and modeling, respectively. Research reported in this publication was supported by the National Institute Of General Medical Sciences of the National Institutes of Health under Award Number F32GM120963 to H.G.Y. and R01GM105984 to A.C.M. The content is solely the responsibility of the authors and does not necessarily represent the official views of the National Institutes of Health.

## Author Contributions

Conceptualization: H.G.Y., A.C.M; Methodology: H.G.Y., P.W.M, J.D., A.C.M.; Investigation, Formal Analysis: H.G.Y.; Writing – original draft: H.G.Y., A.C.M.; Writing-Review & Editing: H.G.Y., P.W.M., J.D., A.C.M; Visualization: H.G.Y., P.W.M.; Supervision: J.D., A.C.M.; Project Administration: J.D., A.C.M.; Funding Acquisition: H.G.Y., J.D., A.C.M.;

## STAR Methods

### CONTACT FOR REAGENT AND RESOURCE SHARING

Further information and requests for resources and reagents should be directed to and will be fulfilled by the Lead Contact, Adam Martin

## METHOD DETAILS

### Fly stocks and genetics

Wild-type embryos with marked membranes and myosin were obtained by crossing *Gap43::mCherry/CyO; sqh::GFP* flies to female virgin *sqh^AX3^;sqh::GFP*. Female progeny not expressing CyO were crossed to OregonR males. The resulting embryos were collected for imaging.

*Sqh-AE* phosphomutant sqh-AE::GFP (sqh-AE) was formed via substitutions of threonine-20 to alanine and serine-21 to glutamate as described in (Vasquez et al., 2014). Sqh-AE mutants were derived from sqh^1^ germline clones carrying sqh-AE::GFP transgene. Mutant/ovoD larve were placed at 37C for 2hr for 3-4 days to create a heat shock using the FLP-DFS method. α-catenin-RNA_i_ embryos virgins of the *shRNA* line, y[1] sc[*] v[1]; P{y[+t7.7] v[+t1.8]=TRiP.HMS00317}attP2 were crossed with a maternal Gal4 driver line, y, w; mat67, sqh::GFP; mat15/CyO, gap43::mCherry-7/TM3. y, w/+; sqh::GFP; mat15/+; gap43::mCherry-7/ P{y[+t7.7] v[+t1.8]=TRiP.HMS00317}attP2 females were mated with siblings and their progeny were examined.

### Image acquisition

Embryos were dechorionated in 50% bleach, rinsed with water, and mounted with embryo glue (double-sided Scotch tape glue dissolved in heptane) ventral side up on a slide. An imaging chamber was constructed with a #1.5 coverslip spacer and was filled with Halocarbon 27 oil (Sigma).

All images excluding the laser ablation experiments were acquired with a 40x/1.2 Apochromat water objective (Zeiss) on a Zeiss LSM 710 confocal microscope. Samples were illuminated with a 488-nm Argon laser and a 561 nm diode laser.

Ablation images were acquired with a Zeiss LSM 710 NLO Laser Scanning Confocal and ablation was carried out with a Coherent Chameleon Ultra II femtosecond pulsed-IR laser and a 40x/1.1 objective.

### mRNA injection

Capped MBS^N300^ mRNA was synthesized using the ~ 900 base pair N terminal sequence of Myosin Binding Subunit fused to a Glycine-Serine linker and an HA tag. A mMESSAGE mMACHINE sP6 Transcription kit (Ambion) was used to generate mRNA. Embryos were dechorionated using same procedure as for live imaging and then desiccated for 5 minutes. The embryos were covered in a mixture of 75% halocarbon 700 and 25% halocarbon 27 (Sigma). The mRNA was injected into the lateral side of the embryo approximately 2.5 hours before ventral furrow formation. The excessive injection oil was wicked off and spacer coverslips (No. 1.5) and a top coverside were added. The imaging chamber was filled with halocarbon 27 oil. Control embryos were injected with 0.1xPBS.

### Image processing and analysis

Images were viewed with ImageJ to select embryos that were the best oriented (http://rsb.info.nih.gov/ij/). A multi-step process was used to project confocal z stacks of the myosin in Matlab (Mathworks). The raw data stack was smoothed in 3D using box smoothing. The approximate location of the embryo’s surface was obtained using the height of the maximum intensity pixels at each position. The resultant image was filtered using bandpass-filtering in the Fourier Domain to create a continuous surface. Myosin signal was averaged for 3 µm above the surface.

Velocity measurements were performed using PIVlab (Thielicke and Stamhuis, 2014) and subsequent analysis was done with custom Matlab code.

The topological tracing of the myosin network was done on the projected 2D image with the *DisPerSE* package (Sousbie, 2010). A persistence cutoff of 5 was used to filter peaks in the signal of topological relevance. *DisPerSE* topologically smoothed the image and outputted the location of all the resultant critical points (local maxima, minima, saddle points) as well as the position of the edges connecting between these critical points. All of the data outputted was at subpixel accuracy. Subsequent data analysis was carried out with customized codes were written in Matlab.

Formally, the *betweenness* centrality for node *u* is defined as 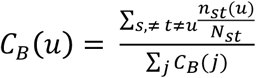, which measures how often node *u* appears on a shortest path between nodes *s* and *t*. *N*_*st*_ is the total number of shortest paths from *s* to *t* and *n*_*st*_(*u*) is the number of these paths that pass through *u*. We then normalize *C*_*B*_(*u*) by the total *betweenness* of all nodes in the network 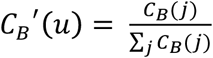. Betweeness of each node was calculated using customized codes written in Matlab employing the built-in function ‘centrality’. Tortuosity of an edge was calculated by dividing the length along an edge by the distance between the edge’s two endpoints. The percent degradation of a network upon myosin phosphatase overexpression was defined by summing the total length of all the edges in the network divided by the average total length of all the edges in the WT data set.

### Computational model

To model the effect of supracellular myosin networks on tissue folding, we employed a continuum elastic shell model, approximating the tissue as an ellipsoid of revolution with major axis *a* and minor axis 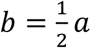. Furrow formation is triggered by modification of the shell’s reference curvature as a function of a prescribed myosin-induced contractility gradient 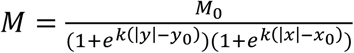 defined on the shell surface, with *x* and *y* the direction parallel and perpendicular to the a-p axis, respectively. In Fig. 6, *x*_0_ = *b*, *k* = 10*b*^-1^, while *y*_0_ = 0.2*b* in Fig. 6B and 0.34*b* in Fig. 6C. This model was adapted from (Heer et al., 2017), and full details of the implementation can be found therein. For our present study, we made one substantial modification to this model by adding an extra term to the energy functional representing the myosin network,

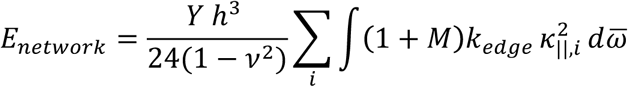

Here the sum is taken over all edges, where 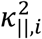 is the surface curvature along the tangent vector parallel to edge *i* at point 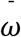 (calculated by taking the projection of the mixed second fundamental tensor along the tangent vector’s direction), *Y* is Young’s modulus, *v* = 0.5 is the Poisson ratio, and *h* = 0.05*b* is shell thickness. The parameter *k*_*edge*_ is a dimensionless multiplier of the stiffness of each edge. In order to best represent the edge topology in a continuum model, we treat this stiffness as a scalar field across the surface given by 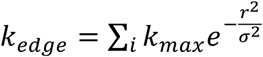, where *r*_*i*_ is the distance of a given point on the surface from edge *i*, and *σ* = 0.0125*b* defines the effective width of each edge. Each edge itself was represented as a straight line with start and end points defined directly from the experimental network trace. Directionally biased networks (Fig. 6B) were constructed by starting with the wild-type network and removing the 50% of the edge most perpendicular to the chosen axis.

## QUANTIFICATION AND STATISTICAL ANALYSIS

Statistical analysis was performed with the MATLAB statistics toolbox. P values were calculated against the null hypothesis using two-sample Kolmogorov–Smirnov test which is a nonparametric hypothesis test that the samples are unequal and from different continuous distributions. All errorbars are +/-standard deviation.

## Supplemental Information Legends

**Supplementary Figure 1:**
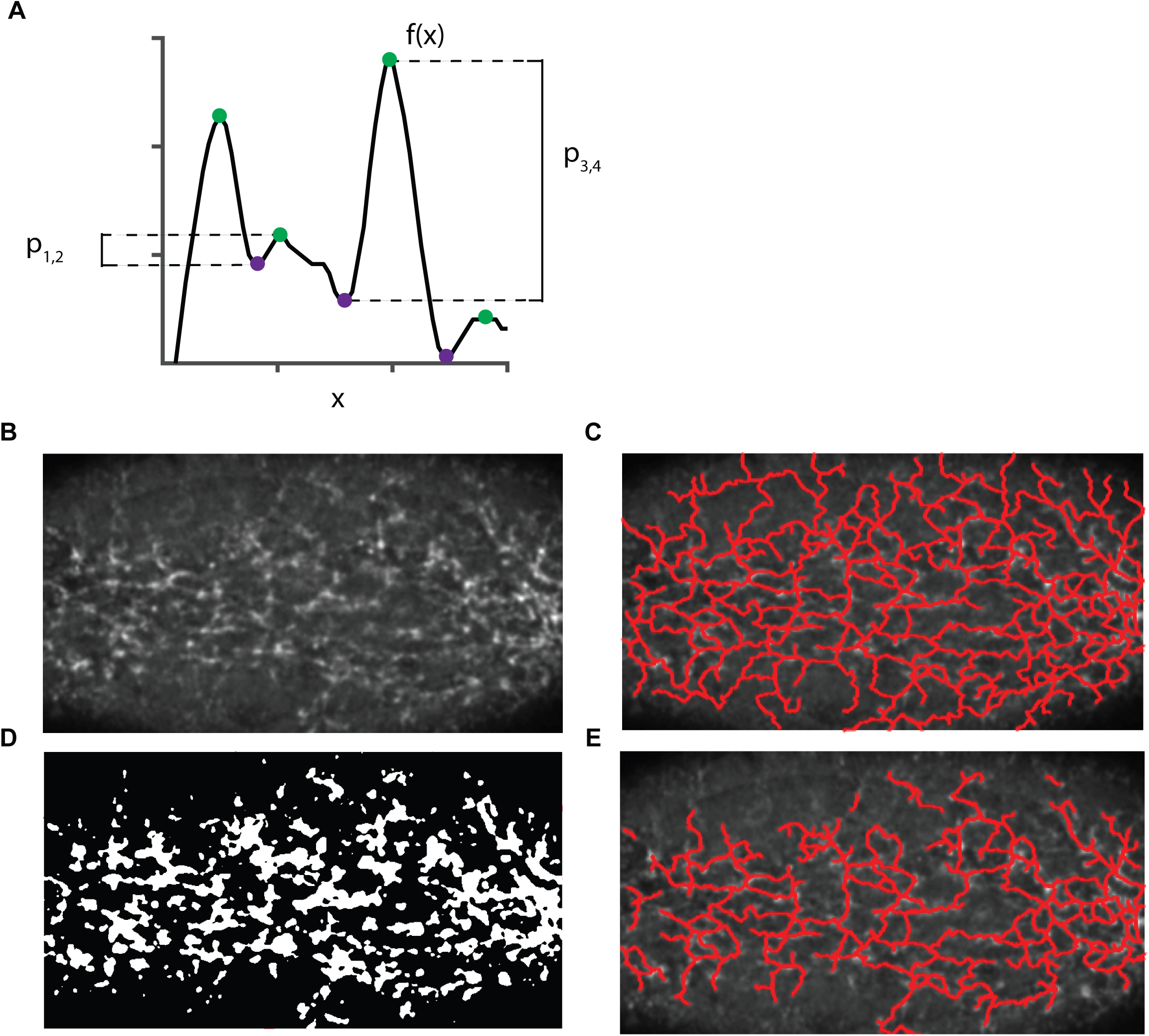
Workflow to trace topologically significant structure in myosin image. (A) For an input function, for example f(x), DisPerSE uses Morse-Smale complex (MSC) analysis to identifying all local maxima (green) and minima (magenta) in an image. Topological relevant extrema are determined by calculating the difference in height or intensity p_ij_ between pairs of extrema (i,j). Pairs above a persistence threshold are maintained and pairs that are below are removed using a topological smoothing. The algorithm outputs the location of the relevant local maxima, and the connections, or integral curves between these peaks through saddle points to neighboring maxima in signal which correspond to lines which trace the top of ridges in the image topographic map. (B) Step 1: A z-stack of myosin around the apical surface of the embryo was projected by retaining pixels near the embryo surface. (C) Step 2: DisPerSE topological tracing was applied which outputs the raw network trace (red) (D) Step 3: Myosin projection was thresholded to create a binary mask identifying the active myosin (as described in (Heer et al., 2017)) from inactive cytoplasmic myosin. The mask corresponded to 0.5*std above the cytoplasmic mean. (E) Step 4: The DisPerSE network trace (red) was filtered to decrease the presence of noise for final trace. Edges that did not overlap with 75% of the mask were discarded. Second, to avoid linking local maxima in distant cells during early pulsing stage of myosin when only a few cells are accumulating apical myosin we removed edges of length greater than ~ 2 diameters or 15 μm.

**Supplementary Figure 2:**
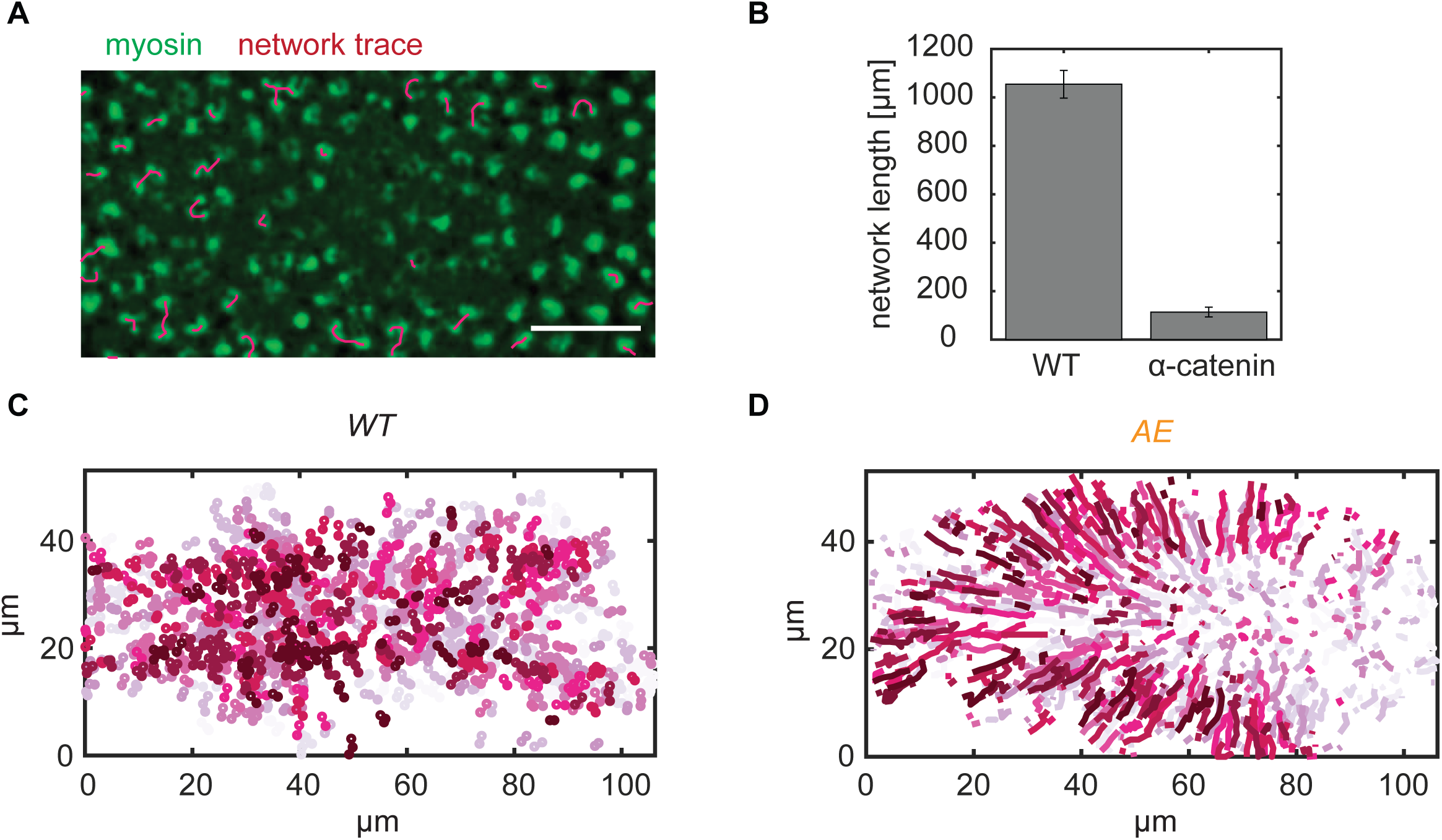
(A) Myosin network trace (red) in an α-catenin-RNA_i_ embryo which has depleted cell-cell junctions. Almost all the network connections are severed. Myosin (Sqh::GFP) is labeled in green. Scale bar: 20 µm. (B) α-catenin-RNA_i_ embryos exhibited a significant reduction of the network length which is defined as the sum of the length of all the edges in the network. (C) Tracks of the nodes in the WT and (D) in the sqh-AE mutant embryo during the 2.5 minutes prior to folding.

**Supplementary Figure 3:**
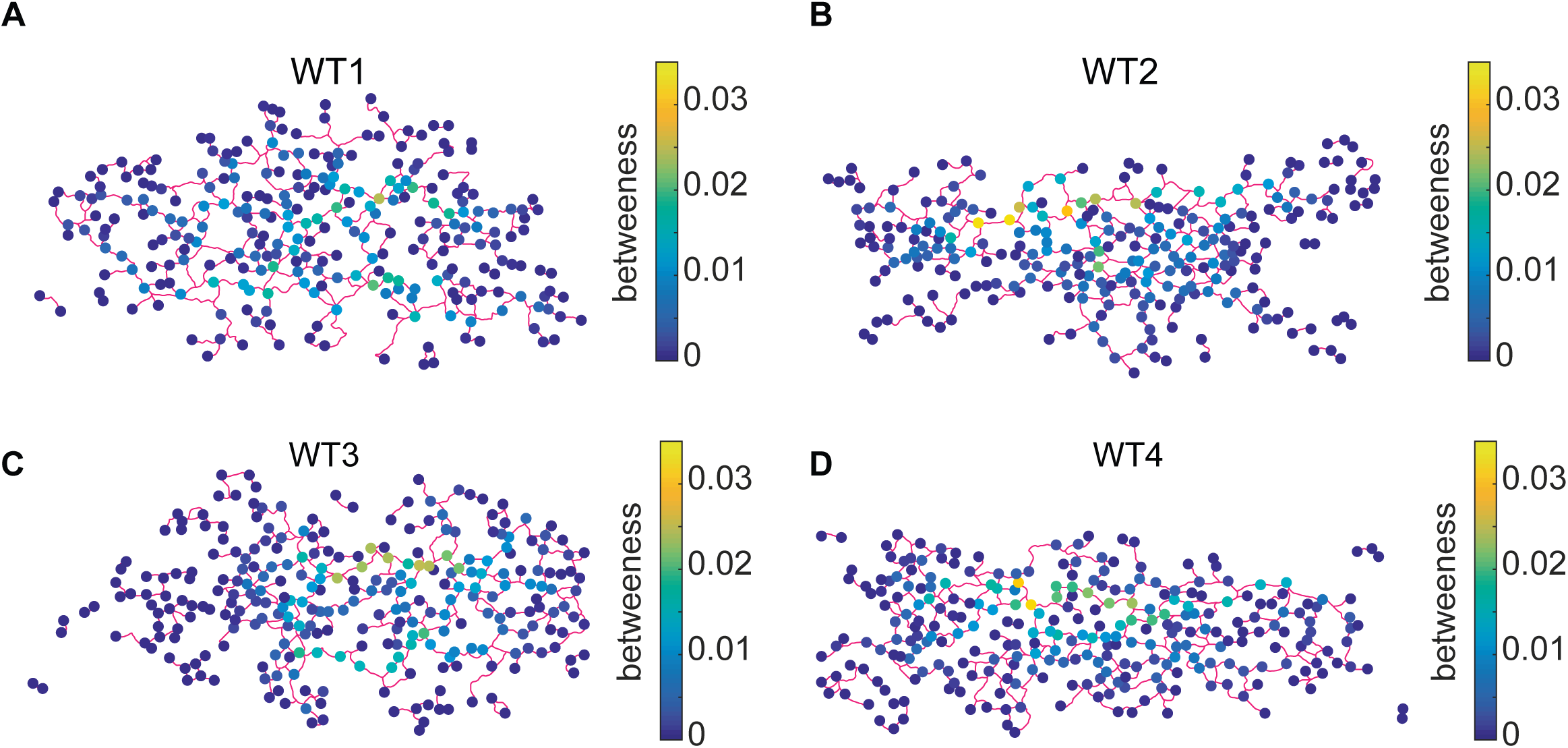
The betweeness centrality calculated for each node in four WT networks at the time of folding initiation (*t=0*).

**Supplementary Figure 4:**
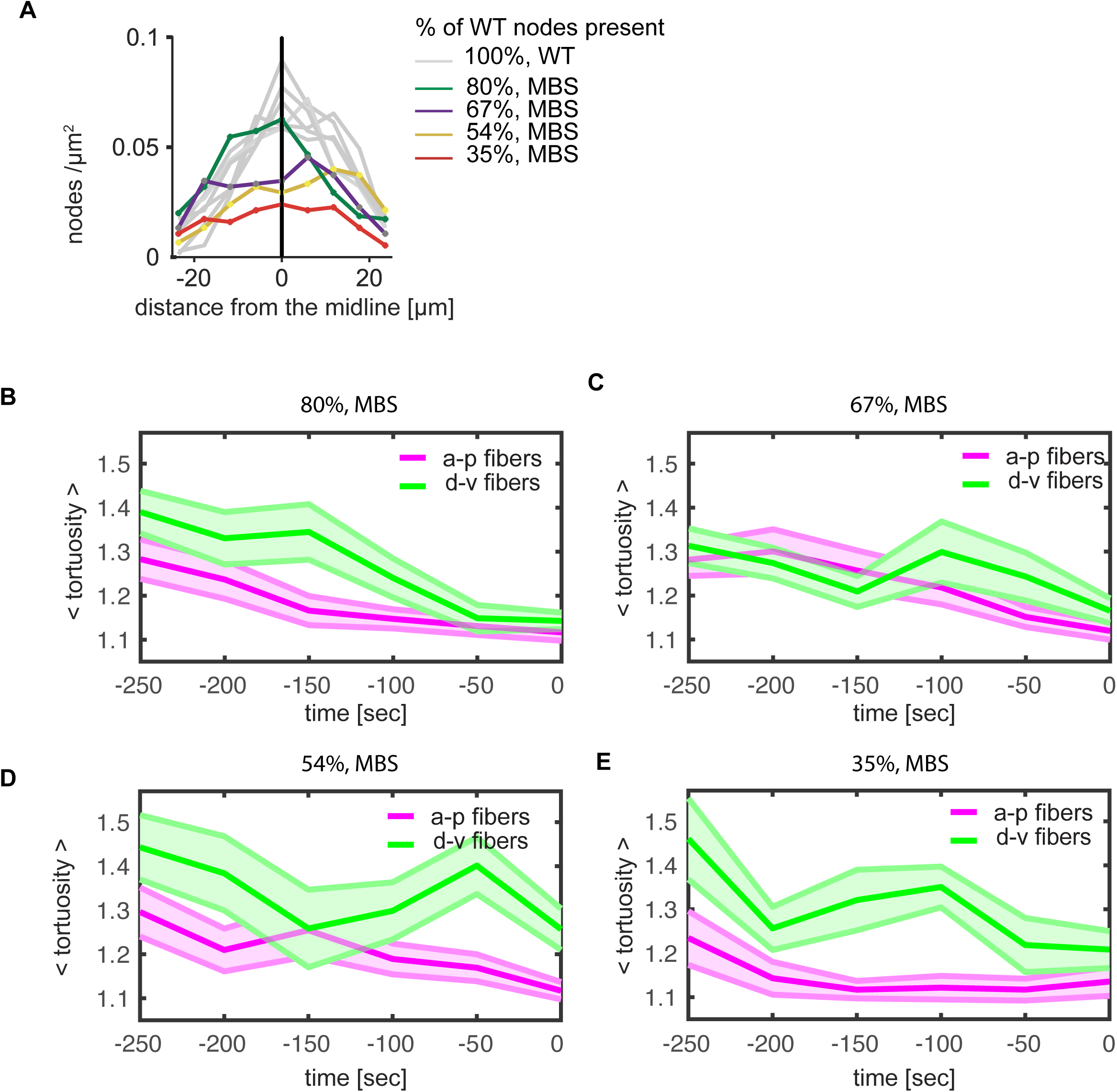
(A) The distribution of nodes from the midline for varying levels of network degradation via myosin phosphatase over expression. The wild-type node density is present in a ventral-dorsal gradient. The gradient is reduced upon myosin degradation. (B) The average edge tortuosity for anterior-posterior fibers (0 – 18 degrees with respect to the midline) and dorsal-ventrally oriented fibers (72 - 90 degrees) for a network that is 80% of the wild-type length. (C) 67% of the wild-type length. (D) 54% wild-type length. (E) 35%. (B-E) each represent a single embryo.

Supplemental Movie 1: Laser ablation of 12 cells. Images are maximum intensity z projections of myosin (Sqh::GFP, green) and cell membrane (Gap43::mCherry, magenta). Scale bar: 20 µm.

Supplemental Movie 2: Myosin network in wild-type embryo. Images are maximum intensity z projections of myosin (Sqh::GFP, green). Scale bar: 20 µm.

Supplemental Movie 3: MBS^CA^ and control injection. Images are maximum intensity z projections of (Sqh::GFP, green). Scale bar: 20 µm.

## References

Barabási, A.-L. (2016). Network science, 1 edn (Cambridge university press).

Baran, P. (1964). On distributed communications: I. Introduction to distributed communications networks (RAND CORP SANTA MONICA CALIF).

Burga, A., Casanueva, M.O., and Lehner, B. (2011). Predicting mutation outcome from early stochastic variation in genetic interaction partners. Nature 480, 250–253.

Chanet, S., Miller, C.J., Vaishnav, E.D., Ermentrout, B., Davidson, L.A., and Martin, A.C. (2017). Actomyosin meshwork mechanosensing enables tissue shape to orient cell force. Nat Commun 8, 15014.

Corson, F. (2010). Fluctuations and Redundancy in Optimal Transport Networks. Physical Review Letters 104, 048703.

Costa, M., Wilson, E.T., and Wieschaus, E. (1994). A putative cell signal encoded by the folded gastrulation gene coordinates cell shape changes during Drosophila gastrulation. Cell 76, 1075–1089.

Davidson, L.A., Ezin, A.M., and Keller, R. (2002). Embryonic wound healing by apical contraction and ingression in Xenopus laevis. Cell Motil Cytoskeleton 53, 163–176.

Dawes-Hoang, R.E., Parmar, K.M., Christiansen, A.E., Phelps, C.B., Brand, A.H., and Wieschaus, E.F. (2005). folded gastrulation, cell shape change and the control of myosin localization. Development 132, 4165–4178.

Derrible, S. (2012). Network Centrality of Metro Systems. PLOS ONE 7, e40575.

Félix, M.-A., and Barkoulas, M.J.N.R.G. (2015). Pervasive robustness in biological systems. 16, 483.

Fernandez-Gonzalez, R., Simoes, S.d.M., Röper, J.-C., Eaton, S., and Zallen, J.A. (2009). Myosin II Dynamics Are Regulated by Tension in Intercalating Cells. Developmental Cell 17, 736–743.

Fernandez-Gonzalez, R., and Zallen, J.A. (2013). Wounded cells drive rapid epidermal repair in the early Drosophila embryo. Mol Biol Cell 24, 3227–3237.

Frankel, N., Davis, G.K., Vargas, D., Wang, S., Payre, F., and Stern, D.L. (2010). Phenotypic robustness conferred by apparently redundant transcriptional enhancers. Nature 466, 490.

Freeman, L.C. (1978). Centrality in social networks conceptual clarification. Social Networks 1, 215–239.

Galea, G.L., Cho, Y.-J., Galea, G., Molè, M.A., Rolo, A., Savery, D., Moulding, D., Culshaw, L.H., Nikolopoulou, E., Greene, N.D.E., et al. (2017). Biomechanical coupling facilitates spinal neural tube closure in mouse embryos. Proceedings of the National Academy of Sciences of the United States of America 114, E5177–E5186.

Guimerà, R., Mossa, S., Turtschi, A., and Amaral, L.A. (2005). The worldwide air transportation network: Anomalous centrality, community structure, and cities' global roles. Proceedings of the National Academy of Sciences of the United States of America 102, 7794–7799.

Gutzman, J.H., Graeden, E.G., Lowery, L.A., Holley, H.S., and Sive, H. (2008). Formation of the zebrafish midbrain-hindbrain boundary constriction requires laminin- dependent basal constriction. Mech Dev 125, 974–983.

Hannezo, E., Dong, B., Recho, P., Joanny, J.F., and Hayashi, S. (2015). Cortical instability drives periodic supracellular actin pattern formation in epithelial tubes. Proceedings of the National Academy of Sciences of the United States of America 112, 8620–8625.

Heer, N.C., Miller, P.W., Chanet, S., Stoop, N., Dunkel, J., and Martin, A.C. (2017). Actomyosin-based tissue folding requires a multicellular myosin gradient. Development, 1876–1886.

Hong, J.W., Hendrix, D.A., and Levine, M.S. (2008). Shadow enhancers as a source of evolutionary novelty. Science 321, 1314.

Hutson, M.S., Tokutake, Y., Chang, M.-S., Bloor, J.W., Venakides, S., Kiehart, D.P., and Edwards, G.S. (2003a). Forces for Morphogenesis Investigated with Laser Microsurgery and Quantitative Modeling. 300, 145–149.

Hutson, M.S., Tokutake, Y., Chang, M.S., Bloor, J.W., Venakides, S., Kiehart, D.P., and Edwards, G.S. (2003b). Forces for morphogenesis investigated with laser microsurgery and quantitative modeling. Science 300, 145–149.

Kermanshah, A., and Derrible, S.J.N.H. (2017). Robustness of road systems to extreme flooding: using elements of GIS, travel demand, and network science. 86, 151–164.

Kiehart, D.P., Crawford, J.M., Aristotelous, A., Venakides, S., and Edwards, G.S. (2017). Cell Sheet Morphogenesis: Dorsal Closure in Drosophila melanogaster as a Model System. Annu Rev Cell Dev Biol 33, 169–202.

Kiehart, D.P., Galbraith, C.G., Edwards, K.A., Rickoll, W.L., and Montague, R.A. (2000). Multiple forces contribute to cell sheet morphogenesis for dorsal closure in Drosophila. J Cell Biol 149, 471–490.

Krueger, D., Tardivo, P., Nguyen, C., and De Renzis, S. (2018). Downregulation of basal myosin-II is required for cell shape changes and tissue invagination. 37, e100170.

Lämmer, S., Gehlsen, B., and Helbing, D. (2006). Scaling laws in the spatial structure of urban road networks. Physica A: Statistical Mechanics and its Applications 363, 89–95.

Leptin, M., and Grunewald, B. (1990). Cell shape changes during gastrulation in Drosophila. Development 110, 73–84.

Martin, A.C., Gelbart, M., Fernandez-Gonzalez, R., Kaschube, M., and Wieschaus, E.F. (2010). Integration of contractile forces during tissue invagination. J Cell Biol 188, 735–749.

Maxwell, J.C. (1864). L. On the calculation of the equilibrium and stiffness of frames. The London, Edinburgh, and Dublin Philosophical Magazine and Journal of Science 27, 294–299.

Monier, B., Pélissier-Monier, A., Brand, A.H., and Sanson, B. (2009). An actomyosin- based barrier inhibits cell mixing at compartmental boundaries in Drosophila embryos. Nature Cell Biology 12, 60.

Nishimura, T., Honda, H., and Takeichi, M. (2012). Planar cell polarity links axes of spatial dynamics in neural-tube closure. Cell 149, 1084–1097.

Parks, S., and Wieschaus, E. (1991). The drosophila gastrulation gene concertina encodes a Gα-like protein. Cell 64, 447–458.

Perry, M.W., Boettiger, A.N., Bothma, J.P., and Levine, M. (2010). Shadow Enhancers Foster Robustness of Drosophila Gastrulation. Current Biology 20, 1562–1567.

Polyakov, O., He, B., Swan, M., Shaevitz, J.W., Kaschube, M., and Wieschaus, E. (2014). Passive mechanical forces control cell-shape change during Drosophila ventral furrow formation. Biophysical journal 107, 998–1010.

Redd, M.J., Cooper, L., Wood, W., Stramer, B., and Martin, P. (2004). Wound healing and inflammation: embryos reveal the way to perfect repair. Philosophical transactions of the Royal Society of London Series B, Biological sciences 359, 777–784.

Rodriguez-Diaz, A., Toyama, Y., Abravanel, D.L., Wiemann, J.M., Wells, A.R., Tulu, U.S., Edwards, G.S., and Kiehart, D.P. (2008). Actomyosin purse strings: renewable resources that make morphogenesis robust and resilient. Hfsp j 2, 220–237.

Rodriguez-Diaz, A., Toyama, Y., Abravanel, D.L., Wiemann, J.M., Wells, A.R., Tulu, U.S., Edwards, G.S., and Kiehart, D.P. (2008). Actomyosin purse strings: Renewable resources that make morphogenesis robust and resilient. HFSP Journal 2, 220–237.

Roper, K. (2013). Supracellular actomyosin assemblies during development. Bioarchitecture 3, 45–49.

Royou, A., Sullivan, W., and Karess, R. (2002). Cortical recruitment of nonmuscle myosin II in early syncytial Drosophila embryos: its role in nuclear axial expansion and its regulation by Cdc2 activity. J Cell Biol 158, 127–137.

Sawyer, J.K., Harris, N.J., Slep, K.C., Gaul, U., and Peifer, M. (2009). The Drosophila afadin homologue Canoe regulates linkage of the actin cytoskeleton to adherens junctions during apical constriction. J Cell Biol 186, 57–73.

Schaffer, C.B., Friedman, B., Nishimura, N., Schroeder, L.F., Tsai, P.S., Ebner, F.F., Lyden, P.D., and Kleinfeld, D. (2006). Two-photon imaging of cortical surface microvessels reveals a robust redistribution in blood flow after vascular occlusion. PLoS Biology 4, e22–e22.

Sharma, A., Licup, A.J., Rens, R., Vahabi, M., Jansen, K.A., Koenderink, G.H., and MacKintosh, F.C. (2016). Strain-driven criticality underlies nonlinear mechanics of fibrous networks. Physical Review E 94, 042407.

Siegal, M.L., and Rushlow, C. (2012). Pausing on the path to robustness. Dev Cell 22, 905–906.

Skoglund, P., Rolo, A., Chen, X., Gumbiner, B.M., and Keller, R. (2008). Convergence and extension at gastrulation require a myosin IIB-dependent cortical actin network. Development 135, 2435–2444.

Sousbie, T. (2011). The persistent cosmic web and its filamentary structure – I. Theory and implementation. Monthly Notices of the Royal Astronomical Society 414, 350–383.

Sousbie, T., Pichon, C., and Kawahara, H. (2011). The persistent cosmic web and its filamentary structure - II. Illustrations. Monthly Notices of the Royal Astronomical Society 414, 384–403.

Sui, L.Y., Alt, S., Weigert, M., Dye, N., Eaton, S., Jug, F., Myers, E.W., Julicher, F., Salbreux, G., and Dahmann, C. (2018). Differential lateral and basal tension drive folding of Drosophila wing discs through two distinct mechanisms. Nature Communications 9, 4620.

Sweeton, D., Parks, S., Costa, M., and Wieschaus, E. (1991). Gastrulation in Drosophila: the formation of the ventral furrow and posterior midgut invaginations. Development 112, 775–789.

Thielicke, W., and Stamhuis, E.J. (2014). PIVlab – Towards User-friendly, Affordable and Accurate Digital Particle Image Velocimetry in MATLAB. Journal of Open Research Software 2, e30.

Umetsu, D., Aigouy, B., Aliee, M., Sui, L., Eaton, S., Jülicher, F., and Dahmann, C. (2014). Local Increases in Mechanical Tension Shape Compartment Boundaries by Biasing Cell Intercalations. Current Biology 24, 1798–1805.

Varner, V.D., and Taber, L.A. (2012). Not just inductive: a crucial mechanical role for the endoderm during heart tube assembly. Development 139, 1680–1690.

Vasquez, C.G., Tworoger, M., and Martin, A.C. (2014). Dynamic myosin phosphorylation regulates contractile pulses and tissue integrity during epithelial morphogenesis. J Cell Biol 206, 435–450.

Whitacre, J.M. (2012). Biological robustness: paradigms, mechanisms, and systems principles. Front Genet 3, 67.

Xie, S., Mason, F.M., and Martin, A.C. (2016). Loss of Gα12/13 exacerbates apical area dependence of actomyosin contractility. Molecular biology of the cell 27, 3526–3536.

Zheng, L., Sepulveda, L.A., Lua, R.C., Lichtarge, O., Golding, I., and Sokac, A.M. (2013). The maternal-to-zygotic transition targets actin to promote robustness during morphogenesis. PLoS Genet 9, e1003901.

